# Dynamics and Selective Remodeling of the DNA Binding Domains of RPA

**DOI:** 10.1101/435636

**Authors:** Nilisha Pokhrel, Colleen C. Caldwell, Elliot I. Corless, Emma A. Tillison, Joseph Tibbs, Nina Jocic, S. M. Ali Tabei, Marc S. Wold, Maria Spies, Edwin Antony

## Abstract

Replication protein A (RPA) coordinates important DNA metabolic events by stabilizing single-strand DNA (ssDNA) intermediates, activating the DNA damage response, and handing off ssDNA to appropriate downstream players. Six DNA binding domains (DBDs) in RPA promote high affinity binding to ssDNA, but also allow RPA displacement by lower affinity proteins. We have made fluorescent versions of RPA and visualized the conformational dynamics of individual DBDs in the context of the full-length protein. We show that both DBD-A and DBD-D rapidly bind to and dissociate from ssDNA, while RPA as a whole remains bound to ssDNA. The recombination mediator protein Rad52 selectively modulates the dynamics of DBD-D. This demonstrates how RPA interacting proteins, with lower ssDNA binding affinity, can access the occluded ssDNA and remodel individual DBDs to replace RPA.

**One Sentence Summary:** The choreography of binding and rearrangement of the individual domains of RPA during homologous recombination is revealed.

## Main Text

In every eukaryotic cell, RPA binds to transiently exposed ssDNA and serves as a hub protein to coordinate essential DNA metabolic processes including replication, recombination, repair, and telomere maintenance ^1,2^. Cellular functions of RPA rely on its high affinity ssDNA binding, its ability to physically interact with over two dozen DNA processing enzymes, and to correctly position these enzymes on complex DNA structures. The precise mechanisms through which RPA functions in many contexts; and more importantly, how it differentiates between multiple DNA metabolic events (DNA replication, repair or recombination) is a long-standing puzzle ^1,3^.

RPA is heterotrimeric, flexible, and modular in structure. It is composed of three subunits: RPA70, RPA32 and RPA14 (Figs.1a, b). The subunits harbor six oligonucleotide/oligosaccharide binding folds (OB-folds; labeled A through F). We refer to the DNA binding OB-folds as DNA binding domains (DBDs; Fig. 1a). RPA binds to ssDNA with high, sub-nanomolar affinity, but can be displaced by DNA binding proteins with much lower DNA binding affinity. Recent studies have suggested that the RPA-ssDNA complex is relatively dynamic ^4,5,6,^ positing a selective dissociative mechanism where not all DBDs are stably bound to the DNA at the same time and microscopic dissociation of individual DNA binding domains occurs.

In all existing models for RPA function, DBDs A and B are assigned as high affinity binding domains. Purified DBD-A, DBD-B and DBD-A/DBD-B constructs bind ssDNA with K_D_ values 2 μM, 20 μM and 50 nM, respectively ^7–9^. The trimerization core made up of DBD-C (RPA70), DBD-D (RPA32) and DBD-E (RPA14) is considered to have a weaker ssDNA binding affinity (K_d_>5μM) ^10^. Additionally, mutational analysis of individual aromatic residues that interact with the ssDNA in either DBD-C or DBD-D show minimal perturbations on ssDNA binding affinity ^6^. Paradoxically, in the recently solved crystal structure of the RPA-ssDNA complex (Fig. 1b), the interactions of all four DBD’s with ssDNA are similar with DBD-C having more contacts with ssDNA bases than DBD-A, DBD-B or DBD-D ^11^. Thus, the exact nature of the contributions from each DBD to RPA function may be more complicated than has been acknowledged and may be influenced by a dynamic component of the DBDs-ssDNA interaction.

Both the N-terminus of RPA70 and the C-terminus of RPA32 interact with a distinct set of RPA-interacting proteins (RIPs) during DNA repair, recombination, and replication. During DNA processing, RIPs must displace RPA from ssDNA. This may be achieved by modulating the DNA binding activity of specific DBDs within RPA. In such a model, a protein that exchanges for RPA does not dissociate all DBDs at once, but individually displaces them after gaining access to DNA that is transiently exposed by dissociation of a DBD. Moreover, if RPA bound on ssDNA were to be considered as a sequential, linear assembly of DBDs as seen in the crystal structure, then depending on the DBD first displaced, a specific DNA binding protein could be positioned at the 5′ or 3′ end of the RPA-occluded ssDNA.

The recombination mediator Rad52 is one example of an RPA-interacting protein. It is a founding member of the Rad52 epistasis group of proteins that orchestrate homologous recombination (HR) and homology directed DNA repair. Specifically, *S. cerevisiae* Rad52 regulates the most critical step in recombination by facilitating replacement of RPA on ssDNA with the Rad51 nucleoprotein filament, an active species in homology search and DNA strand exchange ^12–15^. Nucleation of the Rad51 filament is a slow and tightly controlled process as Rad51 fails to compete for binding to ssDNA with RPA^16^. Rad52 physically interacts with both RPA and Rad51 and assists with the Rad51 filament nucleation. The mechanism by which Rad52 loads Rad51 on the ssDNA is unclear except that Rad51 filament formation is simultaneous with displacement of RPA from ssDNA and likely proceeds through a Rad52-RPA-ssDNA intermediate ^17^. Within this complex, Rad52 was shown to stabilize the RPA-ssDNA interaction^18^, which further mystifies its assigned mechanism of action as a recombination mediator to displace RPA^18^.

To determine how individual DBDs work in the context of the full-length protein and to investigate how proteins such as Rad52 modulate RPA binding, we generated fluorescent forms of RPA containing a non-canonical amino acid (ncAA) that is labeled with the fluorescent dye MB543 by strain-promoted cycloaddition in either DBDs A or D. When positioned near the DNA-binding site, MB543 produces a change in fluorescence signal upon binding to ssDNA. Using direct measurements of the fluorescently-labeled DBD binding to, and dissociating from ssDNA, in the context of the full-length RPA, we show that both DBD-A and DBD-D are highly dynamic, frequently binding to and dissociating from ssDNA. We also show that RPA-ssDNA complexes exist in at least 4 distinct conformational states offering differential access to the ssDNA within this complex. Rad52 interacts with the RPA-ssDNA complex and selectively modulates the dynamics of DBD-D preventing its full engagement to ssDNA and thereby opening the 3′ end of the RPA-occluded sequence.

## Results and Discussion

### Direct read out of DBD dynamics using non-canonical amino acids and fluorescence

Directly monitoring the dynamics (*binding, dissociation or remodeling*) of a single enzyme in multi-protein reactions remains technically challenging. To decipher how the DBDs of RPA function in the context of the heterotrimeric RPA complex, we incorporated MB543, an environmentally sensitive fluorophore to either DBD-A or DBD-D (in RPA70 and 32, respectively, Figs. 1c, d) of *S. cerevisiae* RPA (see methods section for details) ^19^. Both fluorescently labeled RPAs are fully active for ssDNA binding with binding parameters and occluded binding site sizes typical of the wild-type RPA protein (Supplementary note 1 and Fig. S1). RPA labeled at domain A (RPA-DBD-A^MB543^) and domain D (RPA-DBD-D^MB543^) produce enhanced fluorescence upon binding to ssDNA (Figs. 1e, f and Fig. S2) ^19^.

**Figure 1.**
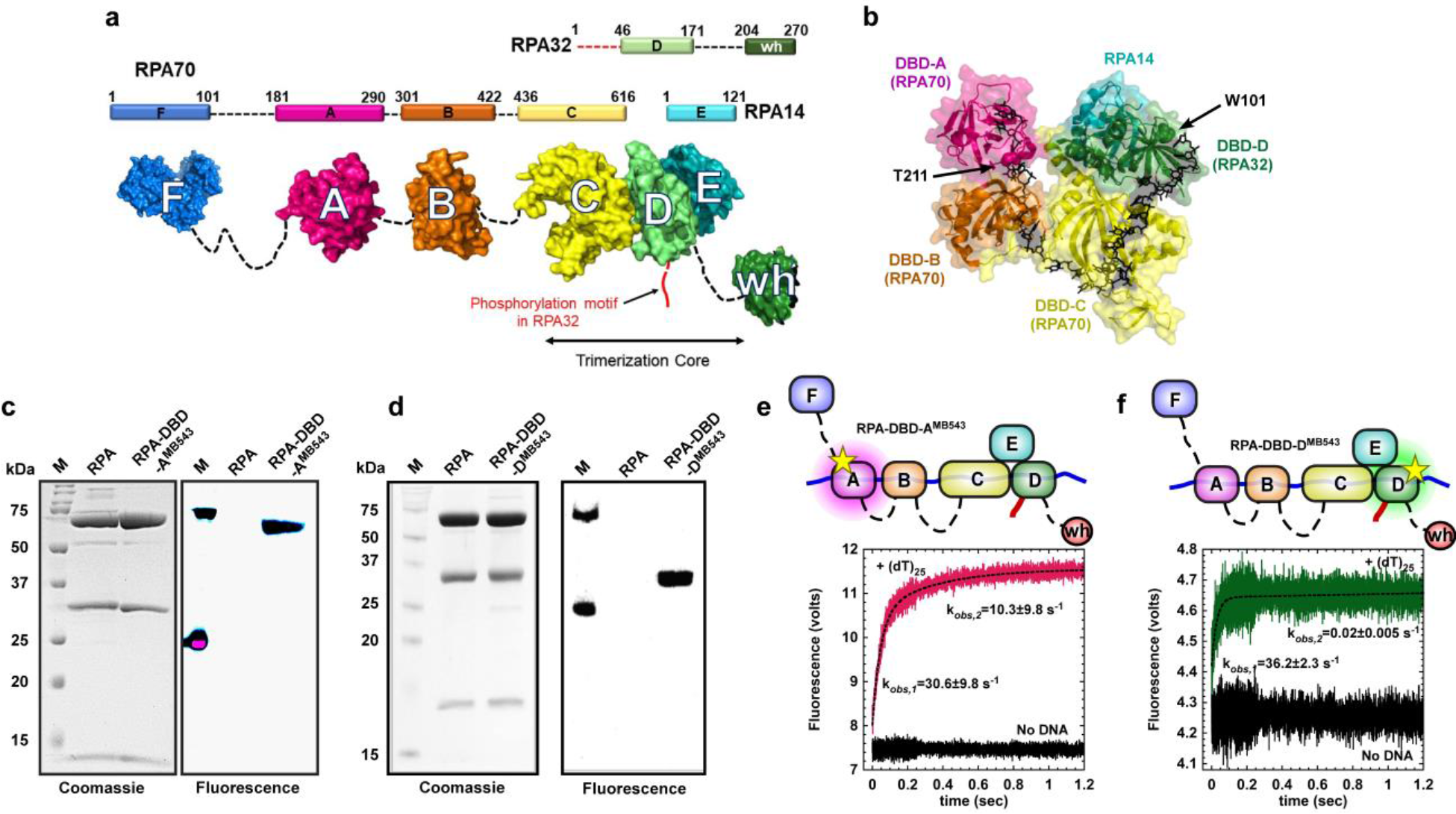
Non-canonical amino acid-based fluorescent RPA report on individual DBD dynamics. **a)** The residue numbers for the three RPA subunits and their respective DNA binding domains (DBDs A-F) are denoted. The winged helix (wh) domain in RPA32 and DBD-F in RPA70 are responsible for interactions with RPA-interacting proteins (RIPs). The N-terminus of RPA32 that gets phosphorylated is shown in red. The crystal structures of the ordered domains are shown as surface representations with intervening disordered linkers as dotted lines (black). DBD-C, DBD-D and RPA14 interact to form the trimerization core. **b)** Crystal structure of the DNA binding domains of *U. maydis* RPA bound to ssDNA (PDB ID:4GNX). Residues T211 in DBD-A and W101 in DBD-D are sites where *4-azidophenylalanine* (4AZP) is incorporated (residue numbering in *Saccharomyces cerevisiae* RPA). The bound ssDNA is shown as sticks (black). **c & d)** Coomassie and fluorescence imaging of RPA complexes labeled with MB543 at either DBD-A or DBD-D. Only the fluorescently-labeled domains are visualized upon fluorescence imaging suggesting site-specific labeling of each domain, respectively. **e & f)** In the stopped flow experiment, RPA-DBD-A^MB543^ and RPA-DBD-D^MB543^ binding to ssDNA were analyzed by monitoring the change in MB543 fluorescence. Robust change in fluorescence depicts engagement of specific DBDs onto ssDNA. Data were fit and analyzed as described in *Methods*.

The ssDNA transactions that involve RPA are a paradigm for reactions where multiple DNA binding enzymes function together on a single DNA template. Knowledge of where, how, and when each enzyme gains access to the DNA in this multi-enzyme milieu is fundamental to deciphering when and how specific DNA repair/recombination processes are orchestrated. Site-specific labeling with MB543 allows us to monitor the dynamics of individual DBDs in the context of the full-length RPA heterotrimer and in multi-protein reactions. Using the fluorescent versions of RPA, we measured the DNA binding kinetics for RPA-DBD-A^MB543^ and RPA-DBD-D^MB543^ providing direct read outs of each domain’s engagement with ssDNA in the context of full-length RPA. RPA-DBD-A^MB543^or RPA-DBD-D^MB543^ were rapidly mixed with ssDNA [(dT)_25_], and the change in fluorescence was measured (Figs. 1e, f). Upon binding to ssDNA, both RPA-DBD-A^MB543^ and RPA-DBD-D^MB543^ produce a change in fluorescence. The data for RPA-DBD-A^MB543^ is best described by a two-step model (k_obs,1_=30.6±9.8 s^−1^ and k_obs,2_ = 10.3±9.8 s^−1^) whereas signal changes associated with RPA-DBD-D^MB543^ fits to a single-step DNA binding model (k_obs_ = 36.2±2.3 s^−1^). The first step in both models is similar and reflects interaction of RPA with ssDNA. The second step for RPA-DBD-A^MB543^ possibly reflects a rearrangement of DBD-A, as has been observed in structural studies ^20,21^. To probe the nature of these differences further, we performed these binding experiments as a function of increasing DNA concentration using a longer ssDNA substrate [(dT)_35_] to ensure that the substrate had ample space for engagement of all the DBDs of RPA (Fig. 2). While measurements of RPA-ssDNA interaction footprints under our buffer conditions yield occluded site sizes of ~ 20 nt/RPA (Fig. S1h), the modularity of the DBDs have been shown to produce occluded site-sizes between 18-28 nt ^22^.

The observed rate for the first association step for both RPA-DBD-A^MB543^ and RPA-DBD-D^MB543^ increases as a function of DNA concentration yielding bimolecular kON values (1.1±0.6 ×10^8^ M^−1^s^−1^ and 2.1±0.4 ×10^8^ M^−1^s^−1^, respectively; Figs. 2a-c, f-h). The second step, observed only for RPA-DBD-A^MB543^, is not linear (Fig. 2c). This is consistent with a conformational rearrangement of DBD-A after the complex with ssDNA has been established and depends on the protein to DNA ratio in the reaction. In these experiments, under conditions where RPA is present in excess over ssDNA, we clearly observe biphasic binding and dissociation/rearrangement phases for RPA-DBD-A^MB543^ (orange and pink traces in Fig. 2b), but not for RPA-DBD-D^MB543^ (Fig. 2g). These data suggest that the dynamics of individual DBDs within RPA differ and may reflect different functional needs; as shown in our single molecule studies (see below).

**Figure 2.**
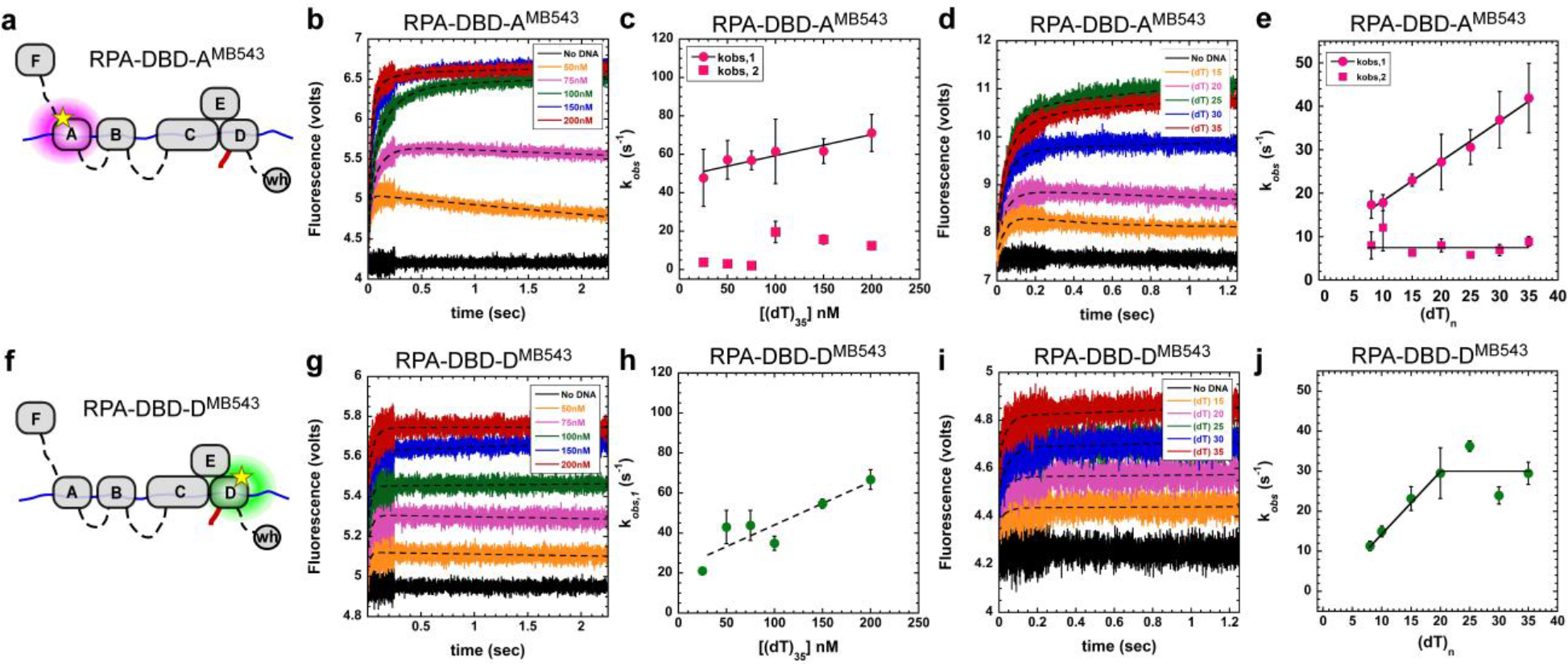
DNA binding dynamics of individual DBDs. **a & f)** Cartoons depicting RPA-DBD-A^MB543^ or RPA-DBD-D^MB543^ binding to ssDNA and producing a change in fluorescence. Stopped flow experiments done with **b & c)** increasing concentrations of [(dT)_35_] ssDNA or **d & e)** with increasing lengths of ssDNA, captures the observed rates in fluorescence change for RPA-DBD-A^MB543^. **g – j)** Similar stopped flow analysis of RPA-DBD-D^MB543^ ssDNA binding dynamics. The data for RPA-DBD-A^MB543^ are best fit using a two-step model whereas the data for RPA-DBD-D^MB543^ fit to a one-step process suggesting distinct DNA context dependent changes in their dynamics.

### FRET analysis confirms primary assessments of DBD-ssDNA dynamics

In our fluorescence intensity experiments, the change in fluorescence arises from environmental changes around the fluorophore upon binding to the ssDNA suggesting that changes in the MB543 fluorescence reflect changes in the electrostatic environment of the dye. (Fig. S3). To better assess whether the dynamics we measure for each fluorescent DBD accurately reflects ssDNA interactions, we used Forster Resonance Energy Transfer (FRET) to capture DBD-ssDNA binding kinetics. RPA binds to ssDNA with specific polarity where DBD-A is positioned closer to the 5’ end of the ssDNA ^11,23^. Similar to the MB543-labeled proteins, we generated Cy5-RPAs where either DBD-A or DBD-D was labeled with Cy5. We next performed FRET experiments with either 5′ or 3′Cy3-end-labeled DNA [(dT)_34_]. On 5′Cy3 DNA, a high FRET signal is observed for RPA-DBD-A^Cy5^, and a medium FRET state is captured for RPA-DBD-D^Cy5^ (Fig. S4a, b). In the corollary experiment with 3’Cy3 DNA, a low FRET state for RPA-DBD-A^Cy5^ and a high FRET state for RPA-DBD-D^Cy5^ are observed (Fig. S4c, d). These experiments are consistent with the expected 5′ to 3′ polarity of RPA binding. Strikingly, the observed rate for the appearance of the RPA-DBD-D^Cy5^ high FRET state (36 ± 2 s^−1^; Fig. S4d) agrees with the rate for change in fluorescence intensity of RPA-DBD-D^MB543^ upon binding to ssDNA (36.2± 2 s^−1^; Fig. 1f). Similarly, the observed rate of 21 ± 1 s^−1^ for the appearance of the RPA-DBD-A^Cy5^ high FRET state (Fig. S4b) is a composite of the two observed phases captured in fluorescence intensity changes of the RPA-DBD-A^MB543^ ssDNA complex (k_obs,1_=30 s^−1^ and k_obs,2_=10 s^−1^; Fig. 1e). The FRET data affirm that the ssDNA binding responsive fluorescence enhancement we observe are a true reflection of specific DBD-ssDNA interactions.

### Differential effects of ssDNA length on DBD conformations

Since each DBD has varying footprints on ssDNA ^11^ we measured the DBD dynamics as a function of ssDNA length and find that the k_obs,1_ increases as a function of ssDNA length for RPA-DBD-A^MB543^ (Fig. 2d, e), whereas the same parameter saturated for RPA-DBD-D^MB543^ at ~20 nt (Fig. 2i, j). On shorter DNA lengths, both binding and dissociation phases are clearly observed for RPA-DBD-A^MB543^ (Fig. 2d: (dT)_15_-orange trace and (dT)_20_-pink trace); however, only a single binding phase for RPA-DBD-D^MB543^ is observed with all ssDNA lengths (Fig. 2i). Since ssDNA and RPA are in molar equivalents (100 nM each) in the experiments with (dT)_n_, the dissociation on shorter DNA probably occurs from intra-subunit competition between the four DBDs of RPA. Due to the different lengths of the flexible linkers between the DBDs, DBDs F, A and B can be considered the conformationally flexible half. In contrast, DBDs C, D and E constitutively interact to form the trimerization core (Fig 1b) and bound to be more conformationally rigid compared to the FAB half. We considered the possibility that the trimerization core might be outcompeting the more dynamic DBD-A (and possibly DBD-B) under conditions of excess RPA or when the length of the DNA is too short to accommodate all the DBDs. To test this scenario, we generated the RPA-FAB fragment containing DBDs F, A and B and labeled it with MB543 in DBD-A (RPA-FAB-A^MB543^). Stopped flow measurement of DNA binding kinetics of RPA-FAB-A^MB543^ yield k_on_ = 1.0±0.1 ×10^8^ M^−1^s^−1^ (Fig. S5d), which is similar to that measured for RPA-DBD-A^MB543^ (1.1±0.6×10^8^ M^−1^s^−1^; Fig. 2b,c), suggesting that DBD-A has intrinsically distinct DNA binding capacity compared to DBD-D, and possibly other DBDs as well. Interestingly, RPA-FAB-A^MB543^ also binds to ssDNA with monophasic kinetics under both conditions of excess protein or shorter DNA lengths (Fig. S5 and supplementary note 2). This suggests competitive binding and rearrangements between the DBDs within full length RPA when the binding sites on ssDNA is limiting. We propose that when a short segment of ssDNA ((dT)_15_ or (dT)_20_) is available, DBD-A rapidly binds and dissociates, whereas DBD-D (and possibly the trimerization core) forms more stable, longer-lived complexes with ssDNA, thus outcompeting DBD-A from short ssDNA substrates. These data also suggest that the interactions of each DBD and resulting conformations of the RPA-ssDNA complex are sensitive to the context of DNA encountered during various DNA metabolic processes in the cell.

### Single molecule analysis reveals the presence of multiple conformational states involving DBD-D and DBD-A

Ensemble stopped-flow experiments described above suggest that the two terminal DBDs of RPA associate with ssDNA with different rates and upon binding to ssDNA RPA commences a complex and dynamic rearrangement of its DBDs. Single molecule total internal reflection microscopy (smTIRFM) was used to directly observe RPA-DBD-A^MB543^ and RPA-DBD-D^MB543^ binding to surface-tethered ssDNA and the microscopic interaction between the labeled domains and ssDNA in the context of the RPA heterotrimer. In the smTIRFM experiments, biotinylated ssDNA (66 nt) was tethered to the surface of the TIRFM flow cell (see material and methods for details). The surface was illuminated with the 532 nm laser, which can excite the MB543 dye when the RPA molecules enter the evanescent field (Fig. 3a and 3b). Binding of a MB543-labeled RPA to surface-tethered ssDNA molecule generates a fluorescence signal in a particular location of the flow cell surface. This signal persists until RPA dissociates, transitions to a dark state and then dissociates, or until the dye bleaches.

Several hundred molecules are observed in the field of view, each yielding a fluorescence trajectory (i.e. change in fluorescence in a particular location on the slide as a function of time) providing quantifiable information on the formation and dissociation of the nucleoprotein complex ^24^. Moreover, environmentally triggered fluorescence changes of the dye in single molecule trajectories can also report on the presence of conformational states in the dye-decorated protein ^25^. The experiments described here were carried out in three stages: first, the surface was observed for 30 seconds to confirm the absence of the non-protein derived fluorescence spots; second, 100 pM RPA-DBD-A^MB543^ or RPA-DBD-D^MB543^ was injected into the flow cell; finally, at 120 seconds protein-containing solution was replaced with the buffer. The last step ensured that the observed changes in fluorescence can be attributed to single RPA molecules. Fluorescence trajectories were extracted from the recorded movies and were normalized using specifically developed Matlab script (described in Methods section and Fig. S10). Only those trajectories that show appearance of the fluorescence signal between 30 and 120 seconds (indicated as ON in Fig. 3c and d) were selected for analysis. Resultant trajectories showed not only appearance and disappearance of fluorescence, but dynamics within the RPA-ssDNA complex (Fig. 3c and 3d). Moreover, transitions between different fluorescence states persisted during the last segment of the experiment suggesting that they truly reflect the conformational dynamics of individual RPA-ssDNA complexes. Global analysis of normalized trajectories for the ssDNA-bound RPA-DBD-A^MB543^ and RPA-DBD-D^MB543^ was performed with ebFRET, a program that determines the number of states using an Empirical Bayesian method to generate Hidden Markov Models ^26,27^. The number of trajectories and states in each experiment are summarized in the Supplemental Table S1 and Fig. S11. This analysis revealed that the fluorescence derived from both proteins best fit a 4-state model, with state 1 corresponding to very low fluorescence and state 2-4 corresponding to increasing fluorescence enhancement. This was true for the segments of the trajectories between 30 and 120 s and for the last 90 s of observation (Fig. 3c and 3d). Segments of the trajectories between 120 and 210 seconds, which can be attributed to the dynamics of a single bound RPA, were used in the quantification of the lifetimes and visitation frequencies for all states.

**Figure 3.**
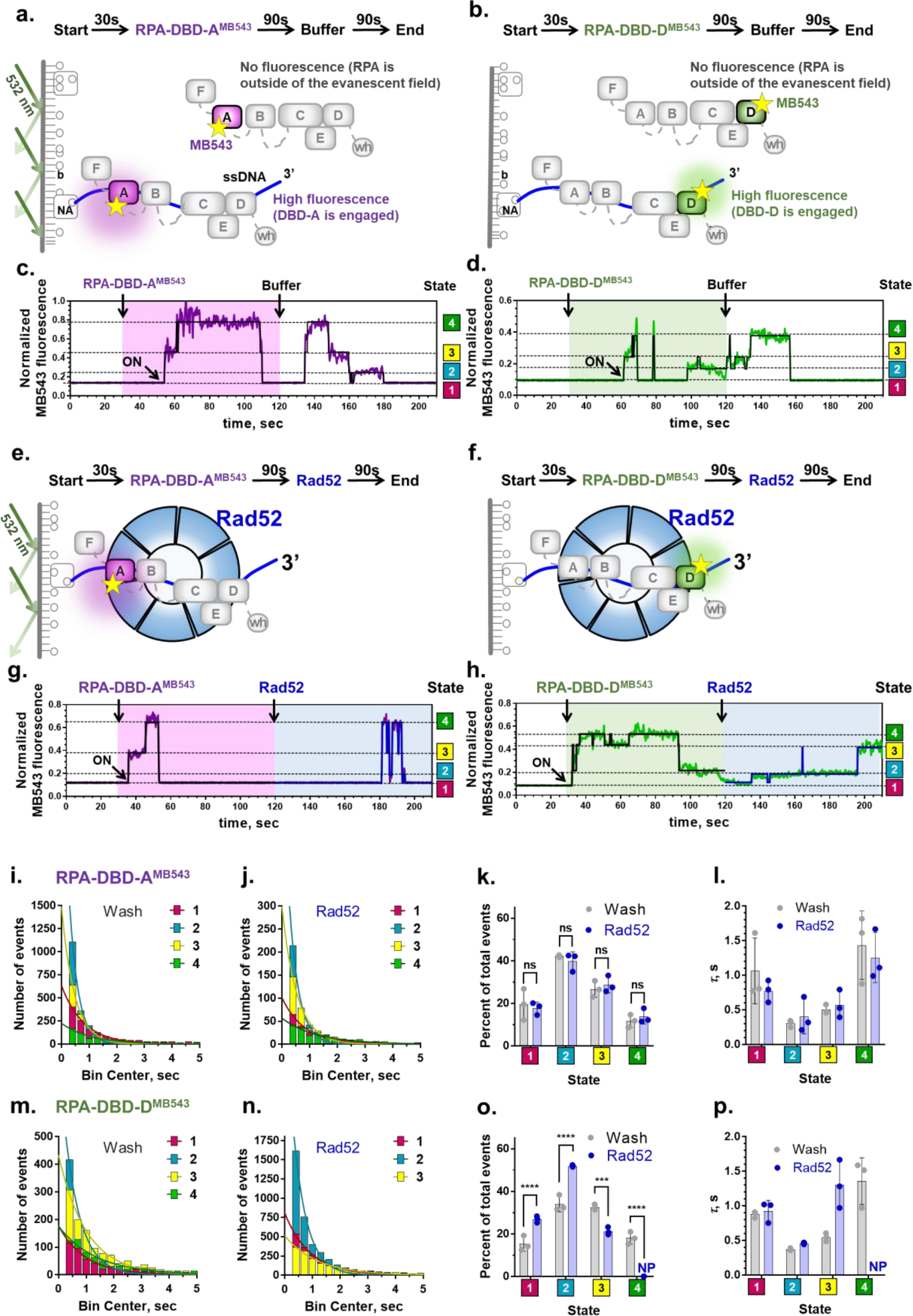
Single-molecule analysis quantifies the conformational dynamics of DBD-A and DBD-D, and the effect of the recombination mediator Rad52 on the accessibility of the 3′-end of the occluded sequence. **a & b)** Experimental scheme for visualizing conformational dynamics of DBD-A and DBD-D, respectively. Binding of fluorescently-labeled RPA (100 pM) to a 66 nt ssDNA (purple line) tethered to the surface of the TIRFM flow cell (grey line) brings the MB543 fluorophore within the evanescent field and its excitation. NA – neutravidin, b – biotin. **c & d)** Representative fluorescence trajectories depicting conformational dynamics of individual RPA molecules labeled within DBD-A and DBD-D, respectively. Purple and green lines represent normalized fluorescence. Black lines represent the results of ebFRET fitting of the experimental data to a four-state model. Additional representative trajectories are shown in Supplemental Figures S6 and S7. The levels for the respective states are indicated by dashed lines. The shaded area on each graph represents the time where free RPA was present in the flow cell. The moment where the RPA molecule binds ssDNA is marked as ON. **e & f)** Experimental scheme for visualization of the effect of Rad52 on the conformational dynamics of DBD-A and DBD-D, respectively. The experimental scheme is identical to that depicted on panels **a** & **b**, except, instead of the buffer wash, RPA in the flow cell was replaced with 700 pM Rad52. In the first 30 seconds of the experiment, and until the appearance of the fluorescence signal (ON), RPA is either not present or is outside of the evanescent field. **g & h)** Representative fluorescence trajectories depicting conformational dynamics of the individual RPA molecules labeled within the DBD-A and DBD-D, respectively. After replacement of RPA in the flow cell with Rad52, the same four conformational states are observed in all RPA-DBD-A^MB543^ trajectories, while RPA-DBD-D^MB543^ trajectories display only three states with the highest fluorescent state (the most engaged state) absent. Additional fluorescent trajectories are shown in Supplemental Figures S8 and S9. **i)** Dwell time histograms for the four fluorescent states obtained by the ebFRET fitting of 258 RPA-DBD-A^MB543^ trajectories from the three independent experiments carried out following the scheme depicted in a. Before fitting, the trajectories were cut from 120 sec (removal of unbound RPA) to 210 sec (the end of the experiment). The dwell times for each state were binned in 300 ms intervals with the center of the first bin set at 400 ms. Each distribution was fitted to a single exponential (solid lines). All the data here and below are summarized in table S1. **j)** Dwell time histograms for the four fluorescence states obtained by the ebFRET fitting of 471 RPA-DBD-A^MB543^ trajectories from the three independent experiments carried out following the scheme depicted in **b**. Before fitting, the trajectories were cut from 120 sec (replacement of unbound RPA with Rad52) to 210 sec (the end of the experiment). **k**) Comparison of the fractional visitation to each state available to RPA-DBD-A^MB543^ alone (grey) and in the presence of Rad52 (blue). Data for each independent experiment is plotted separately. The 2-way ANOVA analysis suggests that there is no significant differences between the visitation frequencies of any of the four states in the presence and absence of Rad52 (p>0.1). **l**) Comparison of the stability of each state available to RPA-DBD-A^MB543^ alone (grey) and in the presence of Rad52 (blue). The data on Y axis are the lifetimes for the respective dwell time distributions. Data for each independent experiment is plotted separately. **m - p**) The same analysis was carried out for RPA-DBD-D^MB543^. Only three fluorescence states were detected in the presence of Rad52. NP – not present. Statistical analysis is performed with ANOVA (***, and **** correspond to p=0.0001 and p<0.0001, respectively).

We attribute state 4 in each case to the RPA conformation where the labeled domain is potentially fully engaging the ssDNA. While it is statistically unlikely that the lowest fluorescence state (state 1) followed by the reappearance of the fluorescence during the last 90 seconds of the experiment is due to the RPA dissociation and rebinding, we substituted the buffer wash with the buffer supplemented with high concentration of ssDNA. In the absence of additional RPA in the solution, ssDNA competitor cannot strip the bound RPA from the DNA, but can sequester all dissociated RPA molecules^5^. As expected, the addition of ssDNA into the reaction chamber had no effect on the RPA fluorescence states (Fig. S12 and Table S2).

To rule out photophysical effects as the source of the MB543 fluorescence states, we repeated the experiments at three different powers of the excitation laser. The lifetimes and the visitation frequencies of the photophysical states, such as blinking, are expected to depend on the excitation laser power, while true conformational states should not display any trend in power dependence^28–30^. Our data summarized in Fig. S13 and Table S3 show that all four states in RPA-DBD-A^MB543^ and RPA-DBD-D^MB543^ have the lifetimes within an experimental error from one laser power to another, thus validating that these states do indeed reflect conformations of the RPA-ssDNA complex. The four fluorescence states and their dwell times were consistent between independent experiments suggesting that the normalization scheme we developed yields reproducible results (Fig. 3k, l, o and p). For both RPA-DBD-A^MB543^ and RPA-DBD-D^MB543^, states 1 and 4 were the most stable with average dwell times around 1 second compared to states 2 and 3 whose average dwell times were between 300 and 500 ms. As evident from the representative trajectories (Fig. 3c, 3d, S6, and S8), RPA spends significant periods of time in the states where DBD-A or DBD-D are not fully engaged thus providing a window of binding opportunity for lower affinity proteins.

In addition, we found that the collective DNA binding affinities of all DBDs produce stable RPA-ssDNA complexes. DBDs A and B have been canonically assigned as responsible for high affinity DNA binding of the RPA complex. By carrying out the single molecule experiments with the RPA-FAB-A^MB543^ we found that it forms a less stable complex on ssDNA and readily dissociates (Fig. S14). These findings agree with results from the bulk stopped-flow experiments where the FAB fragment is exchanged with two orders of magnitude easier than the full-length RPA (Fig. S5 and S15). Fluorescence trajectories recorded for RPA-FAB-A^MB543^ were best fit with the three-state model (Fig. S14b), where state 1 corresponded to free ssDNA and whose lifetime displayed a linear dependence on the RPA-FAB-A^MB543^ concentration (Fig. S14d). Notably, two states, 2 and 3 were present in the bound state of RPA-FAB-A^MB543^ whose two DBDs had been suggested to form a dynamic complex on the ssDNA^31^. The lifetime of the bound state (states 2 and 3 together) of RPA-FAB-A^MB543^ was independent of the RPA-FAB-A^MB543^ concentration, as expected for the ON state of the bound protein. The presence of only two fluorescence states in the RPA-FAB-A^MB543^ construct further confirms that the four states we observed for the full length RPA-DBD-A^MB543^ and RPA-DBD-A^MB543^ are not photophysical states of the MB543 dye.

### Rad52 modulates DBD-D dynamics, but does not affect DBD-A dynamics

To determine the mechanism by which the recombination mediator Rad52 remodels the RPA-ssDNA complex, RPA-DBD-A^MB543^ or RPA-DBD-D^MB543^ bound to the surface-tethered ssDNA in the smTIRFM experiments were challenged with a buffer wash, with or without 700 pM Rad52 (Fig. 3e, f, S8 and S9). The last 90 second portions of the resulting trajectories were normalized and globally analyzed using ebFRET. Dwell times for each state were binned and fit to an exponential decay (Fig. 3 i, j, m, n, Fig. S11 and Table S1). The ebFRET analysis of trajectories for RPA-DBD-A^MB543^ after buffer wash or after Rad52 addition both best fit a 4-state model with the same distribution of states and the same dwell times (Fig. 3 i-l). Trajectories for RPA-DBD-D^MB543^ after buffer wash still fit best to a 4-state model; however, the trajectories collected after Rad52 addition instead best fit a 3-state model (Fig. 3 m-p). Attempts to fit these trajectories with a 4-state model resulted in overfitting and overlapping states. Intensities of the 3-states of RPA-DBD-D^MB543^ after Rad52 addition correspond to the 3 lowest states seen after the buffer wash with the highest state absent when Rad52 was present (Fig. 3 h blue shaded area). RPA-DBD-A^MB543^ visitation frequency of all states remained unchanged between the buffer wash and Rad52 addition (Fig. 3k). In contrast, with RPA-DBD-D^MB543^ occupancy at state 4 is lost with Rad52 addition, state 3 occupancy decreases, while state 1 and 2 occupancy increases (Fig 3o). This suggests that Rad52 selectively modulates conformational dynamics of the RPA-ssDNA complex: reducing the engagement of DBD-D from ssDNA and thus providing access to the 3′ end of the occluded ssDNA.

Formation of the RPA-ssDNA-Rad52 complex depends on the physical interaction between RPA and Rad52, which is mediated by the ssDNA and is confined to the middle region of the Rad52 C-terminal domain^32,33^. To test whether the DBD-D modulation by Rad52 depends on the interaction between the two proteins we used human RAD52 as a control in place of yeast Rad52. Human RAD52 resembles the yeast protein in all its activities but does not engage in the protein-protein interactions with yeast RPA^34^. We found that human RAD52 does not alter the four states of the RPA-DBD-D^MB543^ interaction with DNA (Fig. S16). Thus, the modulation of state 4 is specific for yeast Rad52 and therefore requires Rad52-RPA interaction. To highlight the importance of the interaction between Rad52 and ssDNA for modulating the RPA conformational dynamics we performed another experiment in the presence of a Rad52 inhibitor, epigallocatechin (EGC) (Fig. S17). EGC inhibits the DNA binding activity of human RAD52^35^ and yeast RAD52 (Fig. S17a), but not that of RPA (Fig. S17b). We observed the four fluorescence states of RPA-DBD-D^MB543^ in the presence of Rad52 and EGC (Fig. S17 and table S4). These data suggest that the loss of state 4 also depends on the DNA binding activity of Rad52.

### Model for DBD dynamics and selective modulation within the RPA-DNA-Rad52 complex

Previous models of RPA binding have been based on analysis of subcomplexes or mutational analysis. In contrast, here we have analyzed the properties and dynamics of individual domains in the context of the full RPA complex. This analysis shows that rather than being composed of “high” and “low” affinity domains, the DNA binding domains engage DNA dynamically with DBD-D forming more long-lived complexes with DNA than DBD-A. We also observe interplay between the flexible half of RPA (DBDs-F,A and B) and the trimerization core. Thus, the RPA-ssDNA complex consists of an ensemble of domains that dynamically interact with ssDNA. This suggests that the integration of these interactions result in the diverse functions of RPA. Our data also suggest that the assembly of DBDs on ssDNA are not sequential but rather the result of dynamic, independent interactions between connected DBDs and DNA.

During homologous recombination, RPA forms a complex with the recombination mediator Rad52 ^36^. RPA is in dynamic equilibrium on ssDNA and Rad52 has been shown to increase the residence time of RPA on ssDNA ^18^. Formation of “Early Rad52-bound” RPA and “Late Rad52-bound” RPA are proposed to play distinct roles during Rad51 filament formation and second-strand capture during HR ^18^. The precise interplay between RPA and Rad52 and the nature of their molecular interactions with ssDNA in the complex are poorly understood. The ability to observe in real time the individual RPA DBDs binding to and dissociating from the ssDNA allowed us to build a high resolution mechanistic description of RPA-ssDNA-Rad52 interaction. The heterotrimer of RPA and the heptameric Rad52 ring have similar ssDNA binding sites. Each Rad52 monomer contains an RPA binding site and RPA has two Rad52 binding sites per heterotrimer. Rad52 is believed to interact with the ssDNA backbone, while the DBDs of RPA, especially in the trimerization core engage the bases. Our results show that the stabilization of RPA by Rad52 is a result of both their physical interactions and through contributions of their individual interactions with DNA. We therefore envision a ternary complex where Rad52 and RPA are interacting with one another and are simultaneously bound to the ssDNA molecule. Selective modulation of the DBD-D ssDNA engagement by Rad52 can provide space for Rad52 to interact with ssDNA and stabilize the ternary complex, which now make more extensive contacts with the ssDNA than RPA does on its own. By redistributing the ssDNA between RPA and Rad52, and by reducing the contacts between RPA and ssDNA, such selective remodeling by Rad52 could provide the Rad51 recombinase access to the 3′ end of the RPA-occluded ssDNA, whilst maintaining its interaction with RPA (Fig. 4). Each Rad51 monomer binds to one monomer of Rad52 and to three nucleotides of ssDNA. Six Rad51 monomers are required to achieve a stable nucleation cluster ^37^, which amounts to 18 nucleotides of open ssDNA. This cannot be achieved without the help of a recombination mediator. When Rad52 binds to the ssDNA-bound RPA it modifies the dynamics of the DBD-D domain engagement with ssDNA. Whether this selective modulation extends to the other two DBDs (DBD-C and DBD-E) in the trimerization core remains to be determined. This provides a stretch of ssDNA with sufficient length to initiate the Rad51 filament nucleation. We predict that recombination mediators in other species, including human BRCA2 may operate by a similar mechanism. The details of this mechanism, however, will depend on the intrinsic differences in nucleoprotein filament formation by human RAD51, which nucleates on ssDNA by a dynamic association of RAD51 dimers ^38^ and grows from heterogeneous nuclei ranging in size from dimers to oligomers ^39^.

**Figure 4.**
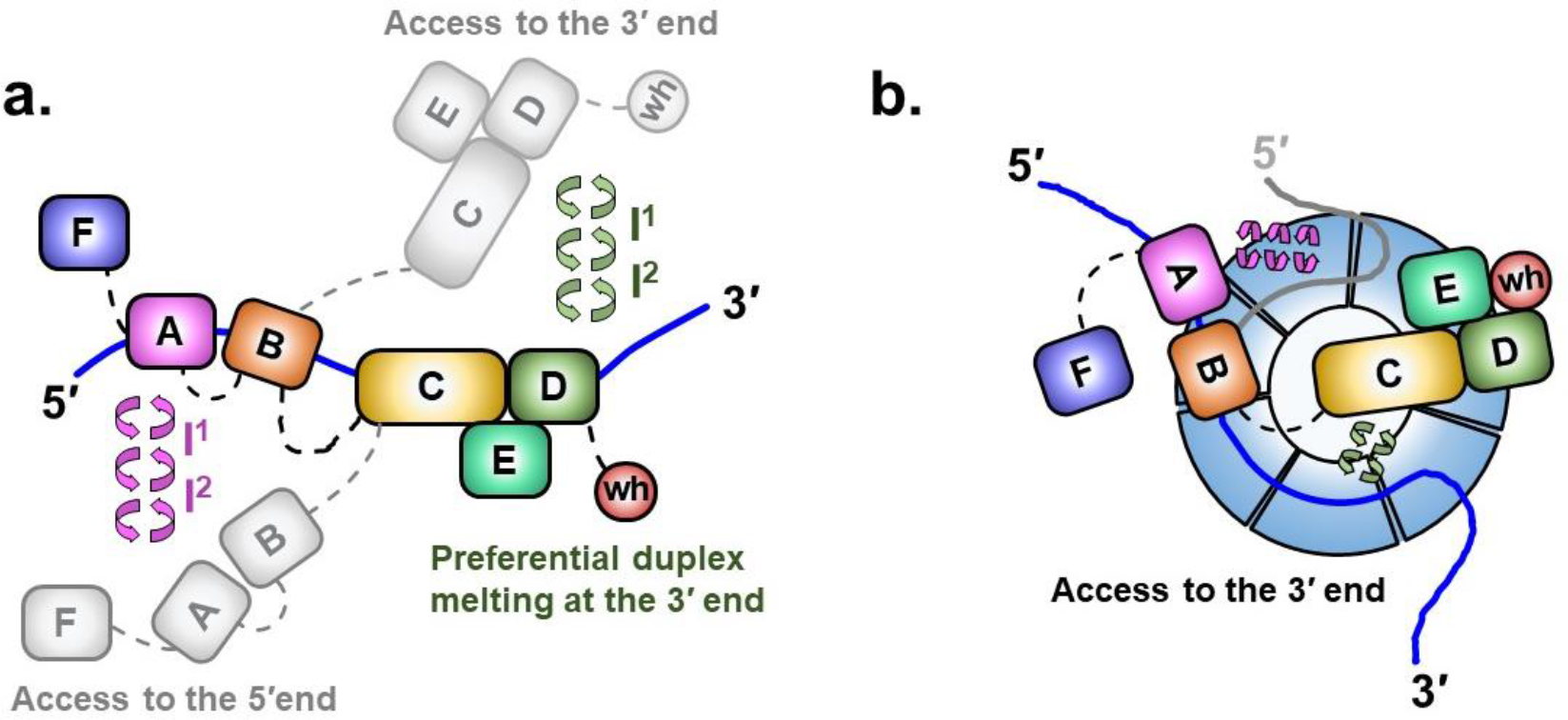
Dynamics of RPA DBDs and modulation by Rad52. **a)** Sequential and directional arrangement of the DBDs allows RPA to occlude 20-30 nt of ssDNA (~20 nt under our experimental conditions; Fig. S1). When RPA is in a stoichiometric complex with ssDNA, or when the ssDNA is in excess, the individual DBDs of RPA exist in a variety of distinct dynamic conformational DNA bound states. Such conformational flexibility allows access to either the 5′ or the 3′ segment of the DNA to other proteins that function in downstream processes. The circular arrows represent the transitions between multiple fluorescence states we observe in the single molecule experiments and which are implied by the bulk stopped-flow experiments. Note that while we illustrate the changes in the conformation of the RPA-ssDNA complex as movement of the DBDs, the same microscopically bound states may arise from ssDNA dissociating and moving away from the respective DBDs. **b)** The DBDs are also selectively modulated by RPA-interacting proteins (RIPs) such as Rad52. In this case, only the DNA binding dynamics of DBD-D, and possibly the trimerization core, is influenced by Rad52. In the ternary RPA-ssDNA-Rad52 complex, the ssDNA is shared between RPA and Rad52, which also interact with one another. The ability of the DBD-D and other RPA elements contacting the ssDNA near the 3′ end of the occluded sequence is constrained. Such selective DBD modulation could promote the loading of the Rad51 protein onto the 3′ end of the ssDNA during homologous recombination.

The myriad cellular roles of RPA in DNA replication, repair, and recombination is also a paradigm for reactions where multiple DNA binding enzymes function together on a single DNA template. Knowledge of where, how, and when each enzyme gains access to the DNA in this multi-enzyme milieu is fundamental to deciphering when and how specific DNA repair/recombination processes are established and utilized. RPA-ssDNA complexes serve as binding targets for the recruitment of appropriate enzymes during DNA replication and various DNA repair processes. Physical interactions between RPA and more than two dozen enzymes have been identified and upon recruitment, the bound ssDNA is handed over from RPA or remodeled in such a way that the DNA is accessible to the incoming enzyme while RPA remains at the site. For example, during nucleotide excision repair, RPA remains at the DNA bubble during most steps in the repair process ^40^. Microscopic binding and dissociation of the RPA DBDs is likely to enable the persistent residence of RPA at the site of repair as well as its ability to coordinate the access to the DNA by helicases and nucleases. Such a mechanism might also be applicable to RPA-like proteins, such as the CST complex associated with telomerase ^41^, that carry a multi-OB fold architecture.

## Acknowledgments

We acknowledge the members in our laboratories for their helpful discussions and suggestions. We thank Dr. Theodore Keppel at the Center for Biomedical Mass Spectrometry Research at the Medical College of Wisconsin for MS analysis.

## Author contributions

N.P., C.C.C., E.I.C., N.J., and E.A.T., performed experiments. J.T., and S.M.A.T., developed the MatLab scripts for data analysis. N.P., C.C.C., E.A., M.S., and M.S.W., conceived and designed experiments and wrote the manuscript.

## Data and materials availability

Constructs used in this study are available upon request. The authors declare that the data supporting the findings of this study are available within the paper and its supplementary information files.

## Supplementary Materials

Supplementary Text

Figures S1 to S18

Tables S1 toS4

Materials and Methods

